# Heterogeny in cages: Age-structure and timing of attractant availability impacts fertile egg production in the black soldier fly, *Hermetia illucens*

**DOI:** 10.1101/2024.08.13.607807

**Authors:** N. B. Lemke, C. Li, A. J. Dickerson, D. A. Salizar, L. N. Rollinson, J. E. Mendoza, C. D. Miranda, S. Crawford, J. K. Tomberlin

## Abstract

Adult behavior is a growing area of interest for those researching the black soldier fly, *Hermetia illucens* (L.) (Diptera: Stratiomyidae), which is affected by underlying demography and spatiotemporal patterns. This greenhouse experiment examined the interaction of age-related effects that can accrue within heterogeneous breeding populations and the potential benefits of delaying an oviposition attractant in concert with restricting mean cohort age. The impetus for this investigation was because if flies are introduced into a mating-cage before old flies are removed or culled, this creates a population of mixed-age and mating-status. We hypothesized this potentially reduces quality among available mate choices, especially in small cages where flies might not be able to spatially segregate. Metrics for fitness included copulation frequency, oviposition frequency, weight of eggs produced, and hatch percentage. “Same”-aged cohorts (maximum 4-d-old at introduction) performed better than highly heterogeneous (1-16-d-old) “mixed” cohorts by mating 2.32-times more frequently and laying 6.58-times more eggs that were 1.17-times more fertile, despite 1.41 fewer observed oviposition events. Delaying the attractant had a significant effect on egg collection weight and led to 1.25-times higher egg yields for same-age populations compared to mixed-aged cohorts where the delaying the attractant had no significant effect. These results are likely due in part to an immediate desire to lay eggs by older females as well as haphazard egg laying in cage material, which was 2.09-times higher for mixed cohorts. The results highlight the importance of constraining the age of breeding populations and removing old adults from cages to improve yields and better manipulate behavior. For those for whom this is logistically unfeasible, providing an attractant box initially and continuously may be the preferred method to trap more eggs from heterogeneous populations.

## Introduction

Energy is a limiting factor for the adult black soldier fly *Hermetia illucens* (L.) (Diptera: Stratiomyidae) (Harjoko *et al*., 2023; Hoffmann, 2021). Because of this, one approach for optimizing reproduction in colony might be to direct the movements of flies more efficiently within regions of breeding environments especially considering both nutritional status (Yuval, 2006; Yuval *et al*., 1998) and spatiotemporal factors (Niyazi *et al*., 2008; Weldon, 2007) are known to dictate reproductive success in other fly species that lek. Speculated methods to achieve this include the manipulation of light, temperature, humidity, and pheromones to lure flies into preferred regions of cages (Salari and De Goede, 2024). Additionally, utilizing physical structures to cordon off mated flies once they have moved into these zones (J. Tomberlin, personal communication), may theoretically reduce intraspecific competition between flies over access to mates (Jones and Tomberlin, 2021). However, to the extent that these are practical are still well into the future in terms of practice and first requires understanding how the various temporal aspects of black soldier fly reproduction interact within a dynamic artificial breeding system to impact fertile egg production.

As construed here, the black soldier fly-breeding system is the group of cage designs and practices employed across the industry that are either copied or derived from the original system proposed in the early 2000s by D.C. Sheppard *et al*., (2002) As the literature reports (both academic and grey), implementations of this breeding system can often be nascent and do not always consider the complexities of evolutionary ecology that underpin insect-mating systems (Shuker and Simmons, 2014). We argue that because flies have a plastic response to their environment (West-Eberhard, 1989), improvements and/or modifications to this system could therefore allow for better alignment of individual flies across the reproductive ‘sequence’ (Giunti *et al*., 2018).

Ideally, these adjustments would lead to an increase in fertile egg production per female, while maintaining or reducing variation, which has historically been cited as being one of the key bottlenecks in reaching economies of scale (Barrett *et al*., 2022; Lemke *et al*., 2023; Pastor *et al*., 2015).

Since at least 2017, companies that mass-rear black soldier flies have bifurcated into those that practice continuous release—which might happen in a single large cage continuously supplied with pupae from adjoining staging areas—or those that implement batch-rearing in a series of isolated cages, a transition that is similar to larval production which likewise utilizes either continuous or batch production methods (Cortes Ortiz *et al*., 2016; Ribeiro *et al*., 2022). These efforts to improve designs and create a more natural habitat are often aligned with the goals of promoting insect welfare and natural behaviors (Barrett *et al*., 2022; Kortsmit *et al*., 2023; van Huis, 2021).

In continuous release, pupae are added into a darkened holding area. As adults emerge from pupal containers, they then join the existing population within the UV-brightened breeding cage, but often prior to the removal of any older cohorts of flies that have also accumulated inside the cage (Dickerson *et al*., 2024). In batch rearing, a similar process of emergence occurs (Dortmans *et al*., 2017); however, populations are isolated from one another, and the cycle can be terminated to clean the cages without disrupting the production of other cages.

From an industrial perspective, this terminology is largely synonymous with continuous and batch systems found in food-processing systems and food plant design (Lopez-Gomez and Barbosa-Canovas, 2005), which makes sense given how much cross-over there is between the two industries’ technologies. However, from a biological perspective, the implementation of what the black soldier fly industry refers to as batch rearing still technically includes the continuous release

of flies because adults emerge from puparia across several days, prior to joining with mating swarms which creates populations with nonsynchronous ages (Dortmans *et al*., 2017) even though the populations from each rearing cycle can be disjunct and thus constitute a batch.

Chiefly, proponents of continuous systems advertise that labor costs and dead time of equipment are reduced (Lopez-Gomez and Barbosa-Canovas, 2005) by having to clean out cages after the termination of every breeding cycle since the total number of cages that need to be serviced is fewer. However, proponents of batch systems claim their method reduces the risk of pest transmission (Manna Insect, 2023) such as red poultry mite, *Dermanyssus gallinae* (Archnida: Mesostigmata), (Mahmoud *et al*., 2023) and other pests and pathogens (Barrett *et al*., 2022; Lemke *et al*., 2023)) while granting the ability to fine-tune parameters during production through integration of remote sensors (Van *et al*., 2022) and computational algorithms (Muinde *et al*., 2023) to autonomously monitor and adjust rearing inputs for optimal outputs in real-time.

Ideally, in a continuous system, daily performance can be estimated as the average over time during peak performance (Lopez-Gomez and Barbosa-Canovas, 2005). However, as alluded to, because not all flies develop or die at the same rates, continuous release of adults into breeding cages (regardless of cage size) can affect the underlying demography of the population. These factors result in highly heterogeneous, mixed-generation cohorts with different mating statuses and physical conditions and probably more so than in a batch system due to overlapping populations of larger sizes. In addition, research has shown a high degree of heterogeny potentially promotes admixture between overlapping generations if employed over long periods, which can contribute to inbreeding depression (Hoffmann, 2021). Such can contribute to collapse, though not necessarily so because not all mass-reared insect populations experience inbreeding depression (Ruiz-Montoya *et al*., 2024), especially with proper management (Cai *et al*., 2022; Facchini *et al*.,

2022). Hence, while there are concerns for the preservation of genetic stock, it is important to note that genetic erosion/deterioration (Bijlsma and Loeschcke, 2012) and the development of domestication syndromes (Lecocq, 2018) have been documented in some, but not all insect species/populations/lines. If such deterioration was pronounced in captive black soldier flies, it could hypothetically lead to a limited ability of black soldier flies to respond to stress, and lower their performance over time compared to wild-type strains (Cáceres *et al*., 2014; Shelly, 2001). Importantly we note that comparing outcomes of artificial selection (intentional or not) applied to the black soldier fly to other mass-reared insects such as those raised for sterile insect technique (Scott *et al*., 2017) is not always apt, considering that the latter must maintain the ability to compete with wild individuals while the former need not.

We predict that inclusion of continuous elements in an insect-breeding system, whether those be through a continuous or batch system (and indeed there are technically “mixed” designs as well) comes at the cost of added variability in egg production due to the heterogeny it propagates. Such was suggested by recent experiments on population structure in which the mixing of two size- (Jones and Tomberlin, 2021) or age- (Dickerson *et al*., 2024) classes led to increased variation in fertile egg production (Dickerson *et al*., 2024; Jones and Tomberlin, 2021). Hence, it follows that heterogeneous populations are likely to have individuals with different mating statuses and physical conditions. In turn, this provides a greater breadth of mate choice and thus increases variation around the mean. However, the practice has yet to be explicitly investigated, nor these predictions. Therefore, this study examines the effect of cohort heterogeny, which is suspected to be greater in continual release systems but still present in batch systems, by comparing fitness between mixed-age cohorts in which flies were allowed to accumulate up to 16 d prior to release, compared to homogenous cohorts with a maximum age of 4-d-old.

Lastly, once cohort heterogeny is reduced, the effect of manipulating the environment to create more enticing lures can be more easily ascertained. Specifically, the larval *(Michishita et al.*, 2023) and microbial (Zheng *et al*., 2013) activity of an inoculated substrate creates a dynamic blend of volatile organic compounds (both local and ambient) (Scieuzo *et al*., 2021) that facilitates fly attraction to egg traps; however, because females are limited by energy reserves acquired as larvae (Lemke *et al*., 2023), they putatively experience a conflict of interest and must allocate time either towards mating or oviposition. By delaying the introduction of the attractant until later in the reproductive cycle, that is when females have possibly, (a) putatively ceased mating, (b) acquired adequate sperm (Malawey et al., 2020; Munsch-Masset et al., 2023), or (c) spent an equivalent amount of time as a wild female might ’flee’ a distant lek to return to the oviposition site (Lemke *et al*., 2023; Tomberlin and Sheppard, 2002). In this study, this duration was equated as 1-d post-peak mating. After the post-mating period (Hull *et al*., 2023; Julita *et al*., 2020; Meneguz *et al*., 2023; Rhode *et al*., 2020), flies can theoretically be provided with an intense signal to lay eggs in traps during the appropriate time of their reproductive ontogeny. In theory, this would also allow them to acquire sperm without the distraction of an attractant, which might otherwise induce them to lay unfertilized eggs in traps (Dickerson *et al*., 2024; Harjoko *et al*., 2023).

Therefore, it is hypothesized that together, (a) young-only cohorts will outperform mixed- aged cohorts, (b) delaying attractant provisioning in synchrony with peak of oviposition during the post-mating period will reduce lead to higher fertile egg production per cage, and (c) this effect will be seen in the young-only cohorts because heterogeny is reduced.

The experiment set-up also allowed the additional investigation as to whether the presence of attractant was inhibitory towards mating due to conflicts of interest between mating and oviposition in energy-limited flies, and whether female attraction towards traps occurred strictly via olfaction or through other means (e.g., vision, learning/memory, artificial selection for traps- seeking behavior over multiple generations).

## Methods

All experiments were conducted in the Forensic Laboratory for Investigative Entomological Sciences (FLIES) Facility (Texas A&M University, College Station, TX, USA) with methods modified from prior studies (Dickerson *et al*., 2024; Jones and Tomberlin, 2021; Laursen *et al*., 2022).

### Larval Rearing

For these experiments, 7-d-old larvae were acquired from EVO Conversions Systems LLC (College Station, TX, USA). Each ’billet’ (Tomberlin *et al*., 2023) of larvae was split in half to create aliquots of ∼5,000, which were then transferred to the center of five plastic storage pans (33.0 × 21.6 × 30.5 cm: l × w × h) (Sterilite® Corp., MA, USA) containing 4 kg of Gainesville diet (Hogsette, 1992) (Producers Cooperative Association, KY, USA) prepared at 70% moisture (Bosch *et al*., 2019) with RO water. An additional 1 L of dry Gainesville diet was spread around the border of each pan to prevent larval escape (Addeo *et al*., 2022; Dickerson *et al*., 2024). Prepared pans were placed inside a walk-in incubator maintained at 16:10 L:D photoperiod, 28 °C, and 27% RH. After 7 d, another 4 kg of Gainesville diet prepared at 70% moisture was provided to each pan. After 10 d, the contents of the pans were sifted by hand using a 6mm sieve (Sona Enterprises®, CA, USA). Pan-level differences that might have arisen during larval rearing were controlled by combining all larvae and prepupae into a single pan after sieving. The wandering motion of the larvae pushed prepupae to the edges of the container, and these were collected. Approximately, 150 g of prepupae were placed in 946 mL cylindrical plastic containers (ULINE® Inc, WI, USA) along with 30 mL of residue (i.e., frass + undigested Gainesville diet) as a bedding material (Dickerson *et al*., 2024). Containers were secured with mesh fabric covers and returned to the walk-in incubator until adults emerged.

### Adult Emergence

A total of four trials were conducted in 2022 lasting from, (a) March 07-13, (b) April 25- May 01, (c) October 06-12, and (d) December 10-16 with larvae reared as described above. The initial two trials had adults that ranged in age from 1-16-d-old to simulate broad age ranges encountered in colonies that practiced continual release. For each trial, 100 males and 100 females were selected to create equivalent age-sex distributions to draw from. To block for the effect of age, the remaining two trials examined responses of 2- to 4-d-old males and 1- to 3-d-old females. Flies that were <1-d-olds (day of eclosion) were not used, because it takes 1 d for the carapace to fully sclerotize (personal observation), during which time flies are likely not sexually receptive (Kobelski *et al*., 2024), as is the case in other insects (Manning, 1967; Ringo, 1996). Emerged adults were identified based on external genital anatomy and placed into 30 x 30 x 30 cm BugDorm 4M3030 insect rearing cages (Bioquip® Products, CA, USA) housed in the walk-in incubator for up to 16 and 4 days; respectively, until all flies (n > 1200 of each sex) had been sorted. While in the walk-in incubator, BugDorms were misted with 35 mL of reverse osmosis (RO) water from a 3.79 L sprayer (Chapin® Manufacturing, NY, USA), twice daily.

**Table 1.** Age ranges for mixed- and same-age treatments.

| Abbreviation | Treatment | Set-up |
| --- | --- | --- |
| Mixed | mixed-age,<br>heterogeneous | 1- to 16-d-old males and 1- to 16-d-old females |
| Same | young-only,<br>homogenous | 2- to 4-d-old males; 1- to 3-d-old females |

### Experiment Design

Experimental units consisted of 12 ‘Insect-a-Hide-pop-up shelters (0.84 × 0.84 × 1.33 m; l × w × h) (Lee Valley® Tools Ltd., ON, CAN), utilized as mating cages. Each mating cage was assigned a replicate randomly (n = 3 replicates x 4 treatments) and arranged in rows of 4, placed 2 inches apart, atop 3 rolling grow tables within an enclosed greenhouse attached to the FLIES facility. Treatments were fully factorial (2 × 2 = 4 levels) and designed to test the effects of and interaction between presence of attractant and delaying availability on mating success, as well as their interaction with cohort age. Half of the mating cages were initially provided with attractant boxes containing inoculated substrate (IA = initial attractant), or boxes absent of substrate (IC = initial control). The remaining attractant boxes, regardless of containing substrate or not, were then delayed until 1-d after peak mating (DA = delayed attractant; DC = delayed control). Including these controls was necessary to rule out the possibility of attraction to the traps via non-olfactory cues (viz. taste).

**Table 2.** Treatment levels and abbreviations for attractant box modification.

| Abbreviation | Treatment | Set-up |
| --- | --- | --- |
| IA | Initial Attractant | Plastic box provided prior to start of experiment, filled with inoculated Gainesville housefly diet as attractant |
| IC | Initial Control | Empty plastic box provided prior to start of experiment |
| DA | Delayed Attractant | Plastic box provided 1-d after peak mating, filled with inoculated Gainesville housefly diet as attractant |
| DC | Delayed Control | Empty plastic box provided 1-d after peak mating |

### Attractant Boxes & Egg Traps

Attractant boxes consisted of plastic shoeboxes (35.6 × 20.3 × 12.4 cm; l × w × h) (Sterlite® Corp., MA, USA) containing 500 g of Gainesville diet prepared at 70% moisture and inoculated with ∼1000 7-d-old black soldier fly larvae (secured from EVO Conversion Systems, LLC, College Station, TX, USA), the presence of which is known to positively influence oviposition (Michishita *et al*., 2023; Zheng *et al*., 2013). Lids were modified to allow volatiles to escape through a 12.7 (L) × 5 (W) cm rectangular slot, which was replaced with nylon screening. Egg traps were constructed by taping together three strips of corrugated cardboard (measuring 10.0 × 3.5 × 1.3 cm; l.0 × w × h). Typically, egg traps were placed perpendicular to the cut-out slot in the attractant box lids; however, at the beginning of the experiment, when ‘delayed’ treatments did not have any attractant boxes present, egg traps were still provided to flies by placing them within the center of the cage floor.

### Observation Schedule

Because mating is stimulated by ultraviolet light (Awal *et al*., 2022; Oonincx *et al*., 2016; Zhang *et al*., 2010), cages were loaded with flies the night prior to the initiation of experimentation under red light (“Day 0”). Observation periods began the following day (“Day 1”), occurring hourly from dawn to dusk, with time rounded to the nearest hour. Observers would walk to cages in a preassigned randomized order, briefly view, and record the total counts of mating pairs and ovipositing flies. Because the duration of mating in black soldier flies is ∼30 min (Manas *et al*., 2023; Masse *et al*., 2022), observed matings were assumed not to have been resampled.

Trials were conducted for six full days and terminated on the seventh. The length of observations was determined by calculating when more than 50% of females had died in a pilot study (unpublished), which was when the cumulative distribution curve for hourly counts reached a horizontal plateau. Each day during the trial, flies were misted with approximately 100 mL RO water at dawn, noon, and dusk.

### Mortality

Each day, between 1200 and 1300 h, dead flies were collected from cages, counted, and sexed. ‘Death’ (or knockdown (Chia *et al*., 2018)) of flies was assessed by gently tapping flies with a size 2 filbert paintbrush (Atool, China). The fly was deemed to have died if there was no response, or if it only gave twitches. This data was not included in any of the analyses presented here.

### Egg Collection

Cardboard egg traps were collected and replaced from mating cages daily between 1200 and 1300 h after dead flies had been collected. Any females that were on traps during the collection were gently nudged so that they would leave. Eggs were then removed from egg traps using a size 2 filbert brush (Atool, China) into a pre-weighed 30-mL plastic SOLO® cup (Solo® Cup Company, IL, USA), and then weighed. As the goal of a breeding program is to promote behaviors that can improve production, no eggs were collected besides those in traps (but refer to (Dickerson *et al*., 2024), for the contrary). All weighing was conducted using an Ohaus Scout® Pro Balance Scale (Ohaus® Corporation, NJ, USA).

### Hatch Rate

To estimate hatch rate, methods adapted from (Dickerson *et al*., 2024)) were used. Each SOLO® cup containing eggs was placed right side up inside a 237 mL screw-top glass canning jar (Choice, China). Jars were secured with a ring lid and a dry paper towel to prevent neonate escape, but still allow some degree of airflow. Jars were placed in a temperature-controlled room set at 26°C, whereupon neonates would crawl out of the SOLO® cup and into the canning jar. After 7 days, the SOLO® cup was then removed from the glass screw-top jar and weighed again so that the difference yielded the mass of neonates that had presumably escaped. Dividing this value into initial egg collection weight yielded ‘hatch rate’ (See discussion section for limitations of this methodology).

**Table 3.** Definition of experimental unit for greenhouse study.

| Variable | Dimensions |
| --- | --- |
| Mating Cage | 0.93 m<sup>3</sup> |
| Initial Density | 100 M:100 F |
| Trial length | 7 days |
| Mean daylight hours | 11.35 ± 2.05 / trial |
| Water | Additional water provided 3 times daily |
| Attractant provisioning | Variable according to treatment |

**Table 4.** Definitions of metrics.

| Variable | Abbreviation | Measured | Units | Dimensions |
| --- | --- | --- | --- | --- |
| Mating events | Mating | For each cage, number of mating events counted at top of hour | Mating events per 200 flies per cage per day | Events/200 flies /0.93m<sup>3</sup>/11.35 mean daylight hours |
| Oviposition in trap | OviTrap | For each cage, number of oviposition events in traps counted at top of hour | Oviposition events in traps per 200 flies cage per day | Events/200 flies /0.93m<sup>3</sup>/11.35 mean daylight hours |
| Oviposition in cage | OviCage | For each cage, number of ovipositing flies in cages (outside of traps) counted at top of hour | Oviposition events in cages per 200 flies per cage per day | Events/200 flies /0.93m<sup>3</sup>/11.35 mean daylight hours |
| Egg collection weight | Egg Weight | For each cage, egg weight refers to all eggs harvested strictly from traps at 12:00 daily and placed in growth chamber for incubation | Total grams of eggs per 200 flies per cage per day | Grams/200 flies/0.93m<sup>3</sup>/day |
| Estimated hatch rate | Hatch | For each egg mass, difference in initial weight and weight after 7 days of incubation, yielded a proxy for weight attributable to hatch, as neonates migrated out of egg collection cups | Percent hatch as the difference between initial egg collection weight and the weight after 7-d of incubation, after which time neonates migrated out of collection cups. | Dimensionless |
| Sex-specific Mortality | FMort, MMort | For each cage, dead flies were removed from cages between 1200-1300 h and sexed. Removed from analyses for simplification. | Sex-specific deaths per 200 flies per cage per day | Deaths/200 flies/0.93m<sup>3</sup>/11.35 mean daylight hours |

#### Statistical tests

Raw data were imported into MS Excel (Version 2302 Build 16130.20218) and later analyzed using R Studio (Version 4.1.3). Prior to statistical analysis, pivot tables were used to generate summary statistics (e.g., means) for fitness metrics subset by treatment and age. For statistical analyses, recorded data for each experimental unit were aggregated, yielding a total of 336 replicate data (12 cages × 7 d × 4 trials = 336). Because of the prevalence of zeroes in the count data, distributions were not normal (as confirmed by Shapiro-Wilk tests) and so all statistical analyses relied on nonparametric tests (viz., Mann-Whitney *U* or Kruskal-Wallis *N*), that examined higher-order interactions before further simplifying models and examining first-order and singular effects. As a simplifying assumption, all reproductive events were assumed to be independent, yet this was not necessarily true, given what is known about polygynandry in captive flies (Hoffmann *et al*., 2021; Jones and Tomberlin, 2021).

### Model Selection

Preliminary modeling that included environmental and mortality data as covariates revealed significant higher-order interactions (i.e., Age × substrate × delay × abiotic × mortality). Because of the complexity of interpreting these nonlinear effects, instead what is presented in the sections that follow are reduced-effect linear models that consider only lower-level interactions (i.e., the effect of treatment (delay × substrate) on fitness or age, while removing the effect of covariates for clarity). This outcome is due in part to flies successive trials experiencing a different suite of temperatures, humidity, cloud cover, radiation, and rainfall.

## Results

### Age x Treatment (Combined Effect)

Iterations of reduced-parameter non-parametric models (i.e., Mann-Whitney *U* or Kruskal-Wallis *N* test) were tabulated to qualitatively analyze which statistically significant results might be under- or overrepresented. In general, treatment effects were not consistently significant (*P* < 0.05) across the entire reproductive chain (i.e., from mating to hatch utilizing the same model architecture), suggesting that delaying the oviposition substrate is unlikely to have a pronounced interaction with age, although some patterns emerged when examining data column-wise across different model architecture.

### Kruskal-Wallis H Test on Subset Data

For example, when subset (i.e., splitting the data set into one group for “mixed” and one for “same”), effects for egg weight were significant and differential in direction depending on cohort which hinted at the presence of an age × treatment effect though this pattern was not universal.

For egg weight, the magnitude of the treatment effect substantially increased with delay for young cohorts, but median values slightly decreased with delay for mixed-aged cohorts, an effect that was not statistically different between mixed- and same-age cohorts (Figure 1).

**Figure 1.**
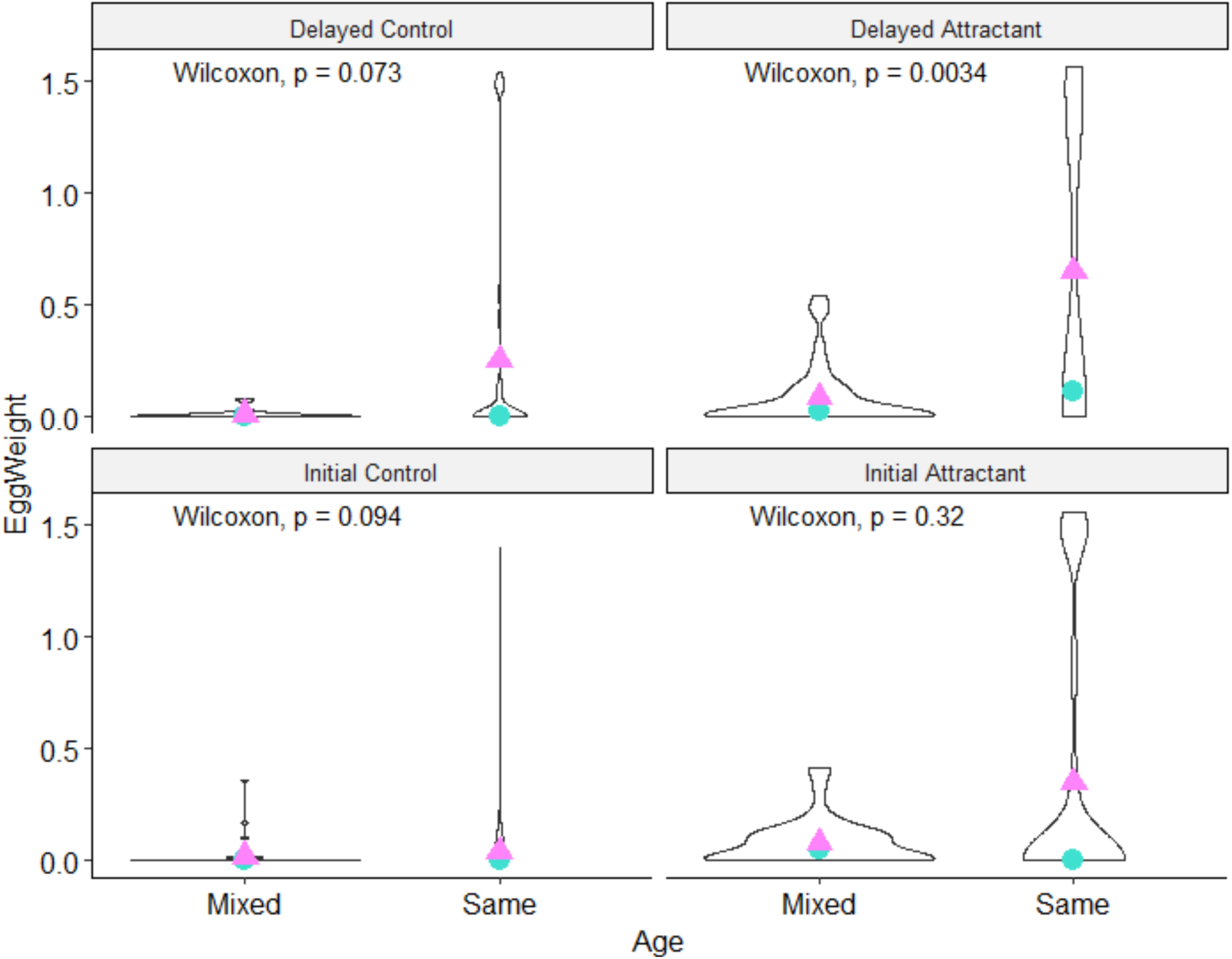
Violin Pot of EggWeight (trapped per cage per day) in response to treatment, zeroes included. Data subset for mixed and same-age cohorts. Blue circles indicate mean values. Pink triangles indicate medians.

In all other instances, one or none of the examined subsets (rather than both same *and* mixed) were globally significant (Table 5). This outcome suggests a need for additional statistical power, or those results that were significant were false positives. However, to truly understand these patterns, nonparametric regressions need to be developed that can adequately model zero-inflated count data (Green, 2021) and are the subject of ongoing work. These models can potentially have better goodness of fit by rectifying nonlinear disparities in abiotic data and providing better separation between groups.

**Table 5.** Kruskal-Wallis *H* Test on fitness data subset by cohort demography. All mating cages had initial populations of 100 M and 100 F flies. ‘Same’-aged flies were 1-3-d old females and 2-4-d old males, while ‘Mixed’- aged flies were 1-16-d old for both sexes. Bolded *P*-values are < 0.05.

| Metric | Subset | Chi-squared | df | *P* |
| --- | --- | --- | --- | --- |
| Mating | Same | 2.67 | 3 | 0.44 |
| Mating | Mixed | 0.887 | 3 | 0.82 |
| OviTrap | Same | 5.52 | 3 | 0.14 |
| **OviTrap** | **Mixed** | **14.7** | **3** | **0.00** |
| OviCage | Same | 2.77 | 3 | 0.43 |
| OviCage | Mixed | 0.860 | 3 | 0.86 |
| **EggWeight** | **Same** | **34.2** | **3** | **1.76e-07** |
| **EggWeight** | **Mixed** | **33.6** | **3** | **2.42e-07** |
| Hatch | Same | 12.3 | 3 | 0.01 |
| Hatch | Mixed | 4.95 | 3 | 0.18 |

### Delay x Substrate

Because the above analyses showed that there was a significant (*P* < 0.05) interaction between age and treatment for at least some of the metrics, the following analyses were conducted to separate treatment effects into their respective components and determine whether the effect was primarily due to delay, substrate alone, or the combined action.

For both statistically significant subsets of EggWeight, Dunn’s test with a Bonferroni correction for multiple comparisons was conducted to determine which treatment groups were significant. After adjusting for *P*-values, the IA-DA (i.e., Attractant × Delay) pair was not statistically significant in the mixed subset (Table 6) but was in the same subset (Table 7). This suggests that the effect of delaying the attractant compared to providing it initially, in terms of improving trapped egg weight, is positive for same-aged cohorts, but there is a null effect towards mixed cohorts.

**Table 6.** Dunn’s post hoc test– pairwise comparisons between different attractant box modifications and their effect on egg weight, subset for ‘Mixed’-aged flies only. These flies were initially 1-16-d old. Zeroes were not truncated. Bolded values indicate *P*-adjusted < 0.05

| | Group 1 | Group 2 | N1 | N2 | Stat | $P$ | $P$-adjusted |
| --- | --- | --- | --- | --- | --- | --- | --- |
| 1 | DC | DA | 42 | 42 | **4.31** | **1.64e-5** | **9.86e-5** |
| 2 | DC | IC | 42 | 42 | -0.663 | 5.07e-1 | 1.00 |
| 3 | DC | IA | 42 | 42 | **3.06** | **2.23e-3** | **1.34e-2** |
| 4 | DA | IC | 42 | 42 | **-4.97** | **6.63e-7** | **3.98e-2** |
| 5 | DA | IA | 42 | 42 | 1.25 | 2.11e-1 | 1.00 |
| 6 | IC | IA | 42 | 42 | **3.72** | **1.99e-4** | **1.19e-3** |

**Table 7.** Dunn’s post hoc test– pairwise comparisons between different attractant box modifications and their effect on egg weight, subset for ‘Same’-aged flies only. Female flies were initially 1-3-d old and males were initially 2- 4-d old. Zeroes were not truncated. Bolded values indicate *P*-adjusted < 0.05.

| | Group 1 | Group 2 | N1 | N2 | Stat | $P$ | $P$-adjusted |
| --- | --- | --- | --- | --- | --- | --- | --- |
| 1 | DC | DA | 42 | 42 | **2.98** | **2.91e-3** | **1.75e-2** |
| 2 | DC | IC | 42 | 42 | **2.13** | **3.30e-2** | **1.98e-1** |
| 3 | DC | IA | 42 | 42 | **5.72** | **1.08e-8** | **6.49e-8** |
| 4 | DA | IC | 42 | 42 | -0.844 | 3.99e-1 | 1.00 |
| 5 | DA | IA | 42 | 42 | **2.75** | **6.13e-3** | **3.68e-2** |
| 6 | IC | IA | 42 | 42 | **3.59** | **3.37e-4** | **2.02e-3** |

In addition, because both combinations that had/did not have attractant (IC-IA, DC-DA) were significant in both subsets, this suggests that the presence of attractant primarily drives oviposition of eggs in traps, while the interaction with delay (IA-DA) is a lone factor in improving egg collection weight by same age cohorts. As expected, there were no differences between controls (DC-IC), and there are significant differences between controls and treatments with attractant (DC-IA, IC-IA).

### Age x Treatment (Full Data Set)

Summary statistics (Table 8) confirm that same x DA combination had the highest mean egg collection weights and hatch rate when zeroes were removed from ‘egg weight’ and hatch rate data (0.89 g/cage/day that hatched with a rate of 60% compared to 0.78 g that hatched 50% of the time for the same x IA combination). This treatment combination had the fewest zeroes (i.e., non-collection events), with 30 cage/day successfully trapping eggs during the experiment. Because the mixed x IA combination was also tied for this value (30 cages/day), this suggests that the IA treatment may be preferred for capturing the most egg-laying events over time for mixed populations, although it will still capture significantly fewer eggs in terms of total weight compared to same x DA combination. Additionally, summary statistics also show that when comparing mixed x IA to mixed x DA combinations, egg collection weight and hatch were slightly higher in the latter when zeroes are removed (though not enough to likely be significantly different), despite fewer events overall (23 cages/day). Oviposition in cage material was strictly lowest in the same x DA combination (2.31/counts/cage/day), followed by the same x IA treatment (2.71/counts/cage/day), and then mixed x IA (4.15 counts/cage/day), and was highest in the mixed x DC combination (12.25/counts/cage/day). Lastly, qualitatively there appears to be little relation between mating counts, and any downstream metric.

**Table 8.**
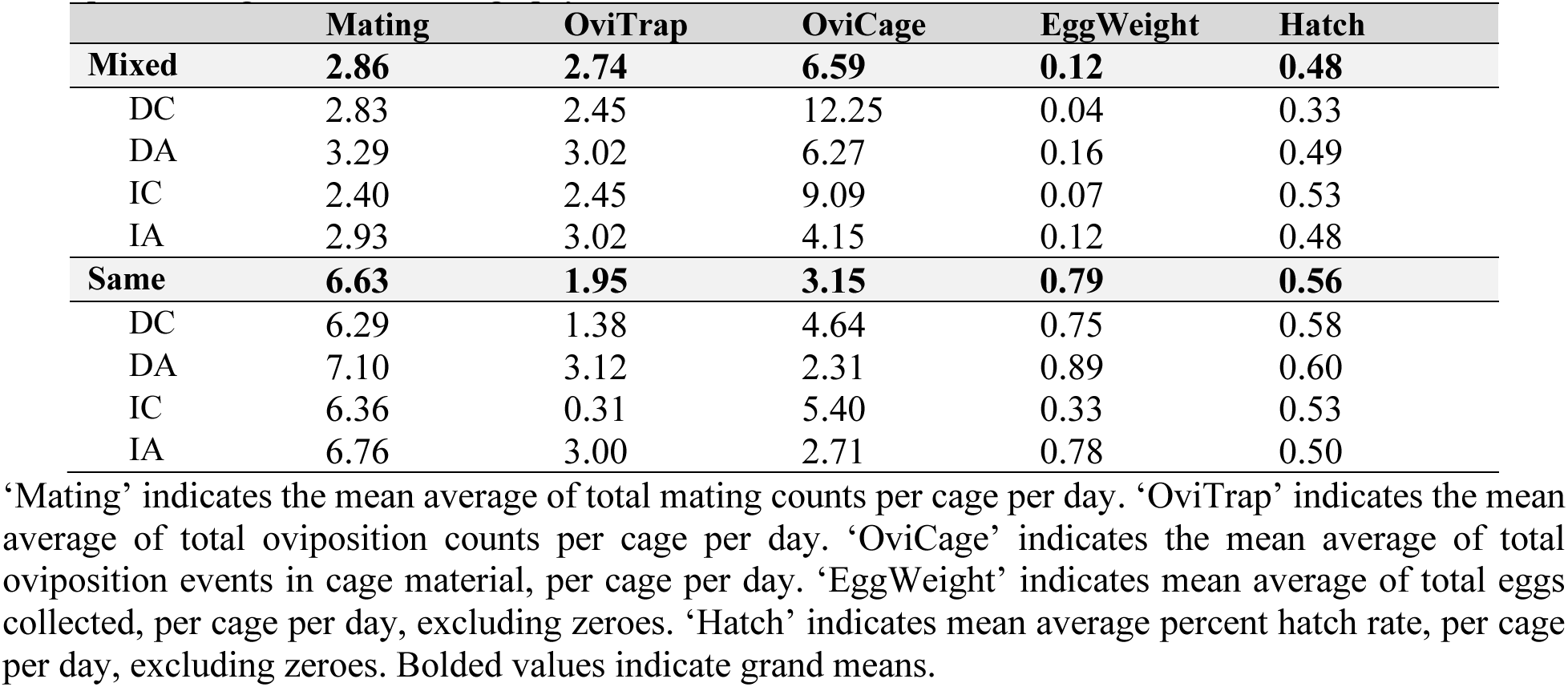
Mean average of fitness metrics (summed per cage, per day) with respect to attractant provisioning and cohort demography.

### Age Primary Effect

Although treatment effects were not ubiquitously significant for all reproductive metrics, Mann-Whitney *U* showed differences between ages that were present across the entire mating chain. Groups were near significant for mating (*P* = 0.06) and would be highly significant (*P* = 0.008) if zeroes were removed (Figure 2). Differences between distributions were also highly significant (*P* = 0.01) for oviposition (Figure 2), nonsignificant (*P* = 0.33) for egg collection weight unless zeroes were removed (*P* = 0.00) (Figure 2), and significant (*P* = 0.02) for hatch (Figure 2), regardless of whether zeroes were removed (*P* = 0.04). Additionally, reasons for their prevalence are given in the discussion.

**Figure 2.**
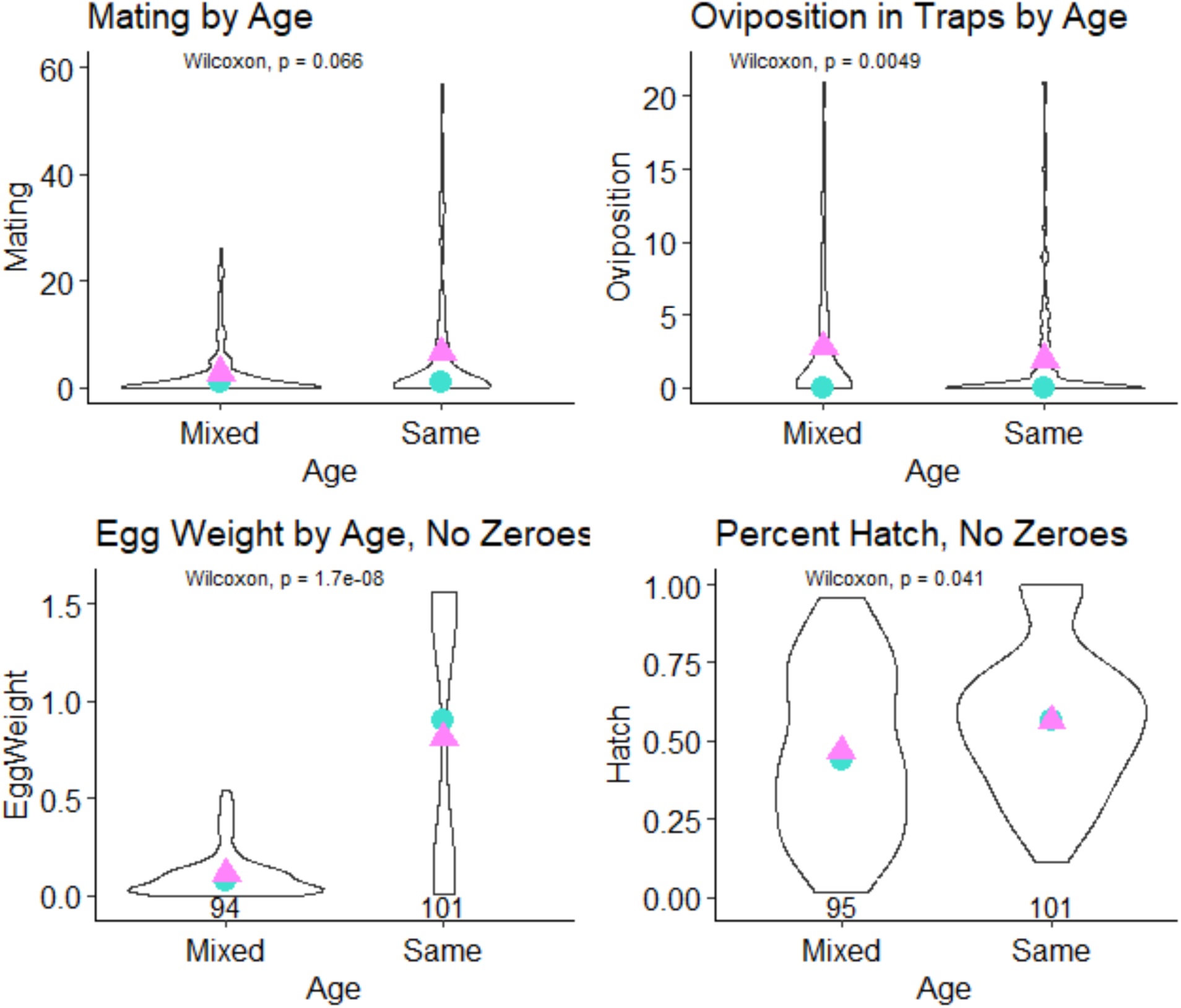
Violin plots for Mating (with zeroes), OviTrap (with zeroes), EggWeight (zeroes removed), and Hatch (zeroes removed) versus cohort demography. Mating and Oviposition given as counts/cage/day. Egg collection weight given as g/cage/day. Hatch given as percent hatch of collected eggs per/cage/day. Blue circle indicates mean values while pink triangles indicate median.

## Discussion

This experiment investigated how modification of oviposition attractant provisioning can differentially affect reproductive behavior in cohorts of adult black soldier flies with either relatively homogenous or extremely heterogeneous age-structure, as measured by mating events, oviposition events, egg collection weights, and hatch rate. Delaying the attractant until 1-day after peak mating was hypothesized as having the best effect at promoting reproduction, and this effect would be more pronounced in cohorts that initially contained flies less than 4-d-old due to a more coherent temporal synchrony with their adult reproductive ontogeny. This is especially relevant to industrial producers, where one of the methods to mass-rear flies is in cohorts of mixed ages. Results from the current study suggest that when an attractant is provided, a given portion of the population in mixed cohorts may not necessarily be at a stage to deposit fertile eggs or otherwise engaged in mating, leading to theorized conflicts of interest and sub-optimal decision-making by flies.

### Treatment Effects

Summary statistics indicate that even in a greenhouse setting, where flies were subjected to a wide variety of environmental conditions during a year, the delay of attractant provisioning can have an obvious effect on some important fitness metrics. Most notably, compared to mixed-age, same-cages that featured an attractant delayed until 1-d after peak-mating expressed 2.16-fold higher mean mating (including all events), similar mean oviposition in traps (including all events), a 2.71-fold reduction in mean oviposition in cages (removing zeroes), and a 1.22-fold increase in mean egg collection weights and highest hatch rates (removing zeroes), suggesting that the act of delaying the provisioning of the attractant can align with underlying reproductive ontogeny during the post-mating period in which females must develop and fertilize eggs. In addition, non-parametric statistics confirm divergent behaviors between age groups with respect to egg collection weight. For both cohorts, the effect of the attractant significantly increased egg collection weights; however, only for young cohorts was there a positive effect of delaying the attractant. This is likely due to poor performance of mixed-aged cohorts, regardless of treatment type, who had much of their oviposition activity diverted to cage material instead.

This result is somewhat counterintuitive, considering the total time flies had inoculated Gainesville available to attract flies towards traps was reduced by 48-h in the delayed treatments. Hence, it follows that as flies age, they should naturally be more inclined to oviposit. The populations with greater mean cohort age at the start of the experiment (i.e., mixed cohorts) were observed attempting oviposition more both in traps and in cages, but this did not lead to greater egg collection weights in traps or hatch rates compared to same-age cohorts. Importantly, egg collection weights and hatch rates of any clutches laid outside of traps were not quantified, as had been done in a prior experiment (Dickerson *et al*., 2024). This is because, from a production standpoint, the relevant quantity is chiefly the eggs that are laid in traps. Certainly, it is possible that performance would have been similar across cohorts if all eggs were included in the total. However, the issue is that collecting eggs outside of the trap would be counter to production optimization by inadvertently promoting the success flies with less discriminant oviposition behavior.

Although prior work showed that populations consisting of a combination of 2-d-old and 4-d-old flies can express multiple (two) peak oviposition periods (Dickerson *et al*., 2024), the present experiment expanded upon this by testing populations that were even more heterogeneous. Because there was no significant difference between providing attractant and initially to mixed-age flies, this suggests a 2-fold nature to the underlying demography (i.e., that some behavioral aspects of the young flies may be dominant in structuring how the population behaves, even though not optimal).

The near ubiquity of non-events (zeroes) in controls confirmed that flies are generally not attracted to cardboard traps atop boxes that did not contain Gainesville attractant, unless perhaps through a random event that eggs were deposited there. This suggests that the primary navigation to traps is olfactory (Zheng *et al*., 2013). Hygroreception of ambient and surface humidity was also thought to be important, as is the case in flies including mosquitoes (Diptera: Culicidae) (Laursen *et al*., 2023) and fruit flies (Diptera: Tephritidae) (Eisemann and Rice, 1989). Once the latter is verified in black soldier flies, this would mean that gravid flies can be lured to traps with in cages through the creation of zones with dramatically higher humidity (Salari and De Goede, 2024).

### Age Structure

Underscoring treatment effects and their interaction with age, large differences in reproductive performance were attributable to age structure alone, with same-aged cohorts performing drastically better than mixed-aged cohorts, confirming our initial hypothesis. Cohorts that had their age restricted experienced several-fold increases in mating, trapped eggs, and hatch rates, despite having fewer oviposition events. Together, this suggests positive knock-on effects to restricting age, as each enhanced aspect of reproduction can benefit those that follow in the chain. Simply put, restricting age created a population of more fit individuals and promoted fertile egg production.

Dickerson *et al*. (2024) explained that reduced and variable performance can potentially be attributed to the presence of old females within breeding populations, while Malawey *et al*. (2020) suggested a root cause in males, namely because senescence can additionally cause black soldier fly sperm quality to deteriorate rapidly. Because of the dynamic interplay of sexually selective forces (Blum, 2012), which engenders the sexes to engage in competition for control over reproductive events (Candolin, 2019); however, neither males nor females can be attributed the blame alone.

Last-male-sperm-precedence has been demonstrated in closely related soldier fly species (Barbosa, 2015) and is typical in fly-mating systems (Shuker and Simmons, 2014). As the name suggests, the sperm of the last male to inseminate the female are favored to fertilize the entirety of her clutch, although in some instances the mechanism may fail if females mate with three or more males (Zeh and Zeh, 1997), leading to multi-sire clutches (Hoffmann, 2021). In the dense environment of black soldier fly industrial cages, ideally, there should be a high proportion of young males in the population so that the chance of an old male being last-to-mate is relatively low. Old males may remove good-quality genes from younger rivals from the females’ genital tract (Shuker and Simmons, 2014), and/or contribute poor-quality genes that have deteriorated through oxidative stress (Missirlis, 2003). Additionally, empirical studies have shown that black soldier fly mating is non-random (Jones and Tomberlin, 2021), especially at lower instantaneous population densities and is governed through multi-step courtship rituals (Giunti *et al*., 2018; Lemke *et al*., 2023). Such rituals involve aerial maneuvers (Tomberlin and Sheppard, 2001), wing displays (Butterworth *et al*., 2021; Meneguz *et al*., 2023), buzzing (Giunti *et al*., 2018), tapping (Giunti *et al., 2018)*, and possibly gentilic stridulation (Eberhard and Gelhaus, 2009). In similar systems, such as lekking fruit flies (Diptera: Tephritidae), old males continue to copulate with females as they age and become more proficient at courting (Papanastasiou *et al*., 2011). Black soldier flies are confined to cages after they mate until they die, even if old flies contribute no genetic material. Together the aforementioned studies explain that the opportunity costs for a population that includes old individuals is relatively high because females could otherwise be copulating with younger mates, and likely contribute to the patterns found during this experiment. While physical methods have not yet been designed to remove old individuals from cages post-breeding, by restricting the heterogeny of the initial starting population’s age, one can at least ensure that the population is of a specified maximum age by the end of a batch-rearing cycle.

Furthermore, by reducing mean cohort age, theory predicts that a reduction in variation around the mean will occur by limiting the breadth of mate choice for the choosy sex(es) (Shuker and Simmons, 2014). This is because in a heterogeneous population, the choosy sex will act on their preferences (Bateson, 1983), and only mate with a select subpopulation. As variation is reduced completely, and limitations (genetic, behavioral, spatiotemporal) are removed, the population reaches an idealized state of panmixia where all individuals freely interbreed and contribute genetic material to the gene pool.

Contrary to our suspicions, reducing variation in underlying age-structure did not reduce variation around the means for fitness, but instead increased it by creating data with abnormally skewed distributions. This finding may be contrary to the findings of a previous study in black soldier flies examining size structure, in which large-only cohorts had less variation compared to small-only and still less than mixed populations (Jones and Tomberlin, 2021); however, we note that the methods of Jones and Tomberlin ((Jones and Tomberlin, 2021)) used standard deviation, which is not appropriate for this study. In our experiment, these outcomes are likely a product of how the data were collected: collecting count data across large stretches of time leads to zero-inflated, skewed distributions that can make describing variation difficult, especially when most conventional statistics (e.g., normal, gaussian, Poisson) assume equal means, medians, and modes (Green, 2021; Perumean-Chaney *et al*., 2013). Because of this, future research should aim to develop methods that can more precisely model variation (Perumean-Chaney *et al*., 2013).

## Conclusion

When examining first- (Treatment × Age) and second order (Attractant × Delay × Age) interactions, in many instances preliminary analyses were not able to differentiate between groups when providing the attractant box with substrate initially or delayed. In some cases, differential behavior was detected between cohort groups, hinting at the possibility that same-age cohorts perform better or the same when the attractant is delayed, while mixed cohorts perform similarly regardless. This suggests young flies need additional time to acquire sperm and lay fertile eggs (in keeping with previous ideas of adult ontogeny (Kobelski *et al*., 2024)), the collection of which can be facilitated by a delayed attractant. Contrarily, old flies, regardless of their mating status, will lay eggs—and not necessarily in traps, since they oviposited in cages approximately twice as frequently—and so to lower the incidence rate of laying in cage material, providing an attractant initially can, (a) direct old flies towards the egg trap, and (b) facilitate laying for younger flies or those that may be introduced successively through a continuous system. Still, logic would dictate that this is less optimal than simply introducing a cohort of young-only flies into the cage, and because same-age cohorts do not statistically perform worse if the attractant is provided initially, logistically this may be the preferred method for producers than delaying the attractant despite a recorded higher performance. Additional modeling of the data will be needed to confirm where economic trade-offs occur as done by Kobelski *et al*. (2024), and future models should consider the spatial and time structure of mating swarms more explicitly. The magnitude of the effect of age on performance seems to qualitatively dominate the interactions, meaning that the effect of modifying attractant timing is smaller than cohort age. As a rule of thumb it is likely that (a) mating is strictly higher in young-aged cohorts; (b) oviposition is strictly higher in mixed populations (unless they are aged beyond the point of being able to lay), and (c) egg collection weight and hatch are higher in young-aged cohorts, irrespective of treatment.

### Limitations

One point to mention is that any reported effect of age is confounded by delayed mating while flies were kept in holding cages prior to their release into experimental cages as an artifact of the experimental design, which in other animal and insect species is known to lower fecundity (Dickerson *et al*., 2024). The reduced egg collection weight of mixed-aged flies may have occurred as a physiological response of flies shunting energy reserves that would normally be used for flight, mating, and an initial bout of oviposition to extend their lives while waiting in holding cages. Reabsorption of oocytes has been previously theorized to occur in black soldier flies (Tomberlin and Sheppard, 2002). Once released into the larger experimental cages, their instinctive behavior putatively shifted to haphazardly laying small clutches of eggs in a last-ditch effort to achieve a small, but net positive fitness (see: Nakamura *et al*. (2016) for females laying additional unviable clutches). Because of this, an obvious recommendation follows that batches of flies should be released into cages within the first 1-2 d of adulthood, or likely suffer from large losses in productivity.

Due to this limitation, which occurred from the desire to keep methods consistent with past work (and build a series of experiments for a future meta-analysis), there is an important nuance to stress that the heterogeny being replicated in this experiment does not directly mimic what occurs at industrial scale at the beginning of a rearing cycle. This is because industrial producers likely are not storing flies for extended periods of time prior to release, unless by accident. Instead, this experiment should be thought of as giving clues to some interactions that happen as cohorts overlap and become increasingly heterogeneous populations, as well as documenting behaviors that occur in sub-optimal caged environments.

In addition, any findings may be confounded by spatial effects. Although each cage within the experimental array was re-assigned treatments randomly at the beginning of each trial, certain cages were situated closer to a wet wall within the greenhouse. Because of this, reproductive behavior was likely impacted along a gradient of increasing humidity, that interacted with weather and climate. Specifically, casual observations throughout the experiments (leading to a heuristic) revealed that cages that were situated furthest from the wet wall had the highest incidence of oviposition in cage material, primarily on their wet-wall-facing edge, followed by sides that neighbored cages with high amounts of oviposition in the walls. This suggested that where black soldier flies oviposit might be mediated through zones of local humidity (although past research has suggested a preference for dry areas (Booth and Sheppard, 1984)), as well as volatiles emanating from conspecific eggs (Zhang *et al*., 2023) and/or larvae (Zheng *et al*., 2013). We speculate this can create a positive feedback loop, inducing additional females to lay outside of traps. This finding is in keeping with what industry producers have communicated(i.e., that local humidity is important for directing flies towards egg traps) (Salari and De Goede, 2024), and adds to a current report that there are contrarily no known dynamic effects of ambient humidity (Kobelski *et al*., 2024).

Lastly, the reported hatch rate is not a measure of true hatch, since the methodology invertedly placed selection pressure on larvae that escaped the SOLO® cup, which not all larvae did. In addition, some of the weight loss between the initial and 7-d weighing events can putatively be attributed to desiccation over the 7-d incubation period. The measure also ignores the fact that hard chorion is left behind after hatching and that some eggs were nonviable. The difference in these respective quantities likely varied across replicates (collections per cage per day) (Dickerson *et al*., 2024) and cannot be accurately distinguished without examining eggs individually, which is incredibly time-intensive (Dickerson *et al*., 2024; Mossman *et al*., 2019). For logistics’ sake an estimation was preferred, and so needs to be viewed as such. It is also possible that cup orientation (i.e., placed vertically or sideways) negatively affected the hatch rate, since larvae at minimum might have to travel twice as far to leave the cup (i.e., to the top of the rim, and then along the outside surface of the SOLO® cup if placed in a vertical orientation; versus just to the edge and onto the bottom interior of the canning jar placed in a horizontal orientation), although this is unverified. However, since all replicates were handled identically, such an effect would have been experienced uniformly. Future experimental work might seek to improve upon the methodology by collecting training data on rates of weight loss due to desiccation to parameterize a model, or by examining how effects of scale on hatch rate (since, several kg of eggs are collected per day, at scale). In addition, computer vision might be leveraged to increase the precision of hatch rate estimates, such as by modifying the *HistoNET* program (Sharma *et al*., 2020) that is used by the industry for automated egg counting (Dickerson *et al*., 2024).

## Acknowledgments

We would like to thank our anonymous reviewers for helping revise the manuscript and acknowledge Drs. Jonathan Cammack, Jessica Yorzinski, Seppe Salari and Lotte Joosten, for their interest and critiques, as well as additional contributions from Sienna McPeek, Rachel McNeal, and Drs. Jennifer Rhinesmith-Caranza and Casey Flint.

## Funding

This material is based upon work supported by the National Science Foundation Graduate Research Fellowship under Grant No. 1746932. Any opinion, findings, and conclusions or recommendations expressed in this material are those of the authors and do not necessarily reflect the views of the National Science Foundation.

In addition, this research used resources from the Texas A&M AgriLife Institute for Advancing Health Through Agriculture.

